# Interactions of Interaural Time and Level Differences in Spatial Hearing with Cochlear Implants

**DOI:** 10.1101/2025.01.29.635454

**Authors:** Sarah Buchholz, Susan Arndt, Jan W. H. Schnupp, Nicole Rosskothen-Kuhl

## Abstract

Normally hearing humans can localize sound sources quite accurately, with minimum audible angles as small as 1°. To achieve this, our auditory pathways combine information from multiple acoustic cues, including interaural time and interaural level differences (ITDs and ILDs). Patients relying on cochlear implants (CIs) to hear the world cannot match normal performance. These deficits are most pronounced in patients with little or no hearing experience early in life, and they appear to result from an impaired sensitivity to ITDs, but not to ILDs. However, little is known about how ITD and ILD sensitivities develop and interact in an early deafened auditory system shortly after CI implantation. We fitted neonatally deafened rats with bilateral CIs, and, providing informative ITDs and ILDs from stimulation onset, trained them to lateralize CI stimuli. These animals were exquisitely sensitive to both ILDs and ITDs of CI stimulus pulses, and combined information from both cues in a weighted sum. Importantly, ITDs are weighted heavily in our CI rats, such that only very modest ITDs pointing in one direction can confound quite large ILDs pointing in the opposite direction. This underlines the importance of informative ITDs for maximizing the potential for spatial hearing with CI devices.

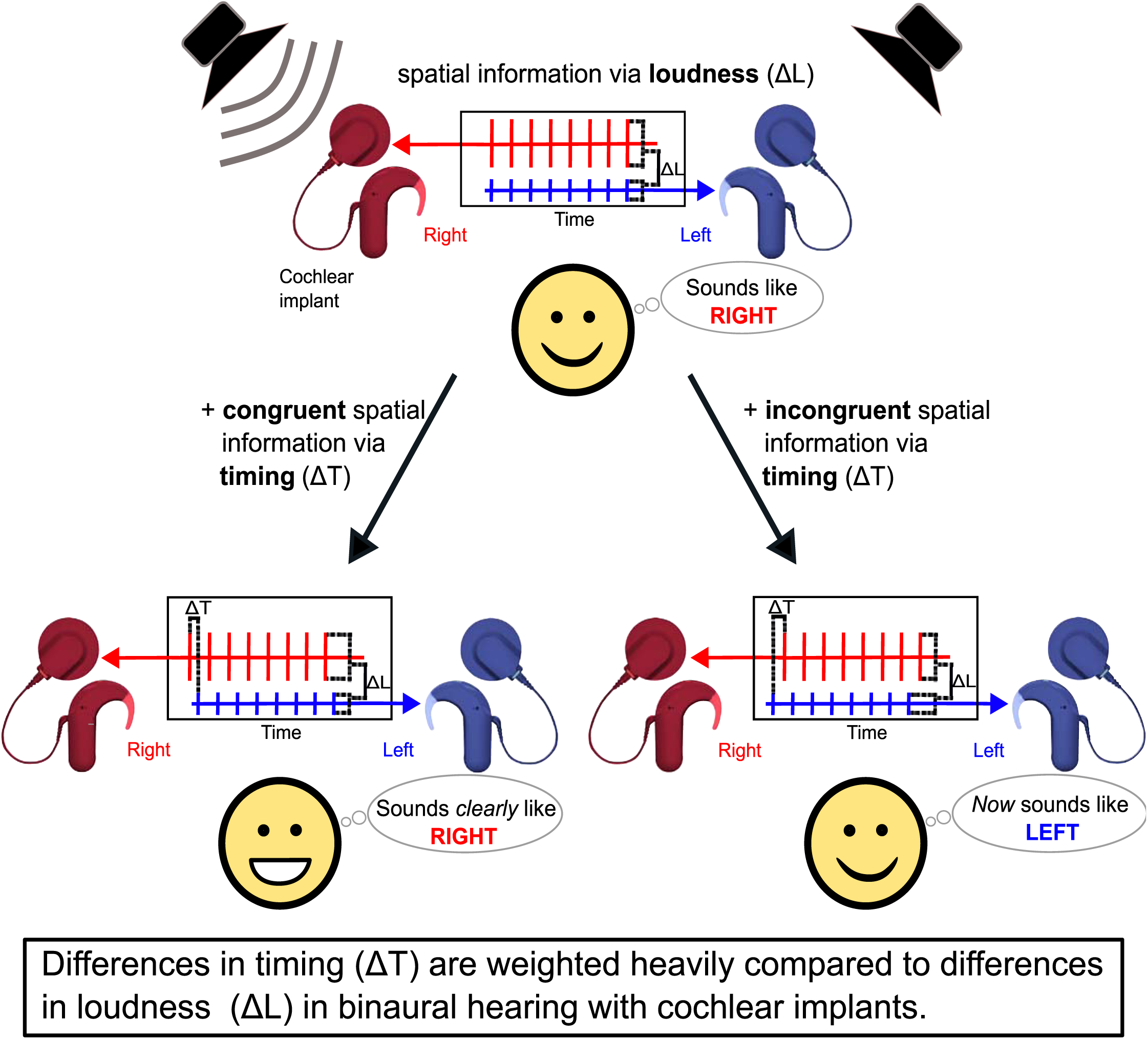

## Introduction

Our ability to hear, to detect, identify and also to localize sound sources, allows us to interact with and navigate through our acoustic environment. Spatial hearing relies on the brain’s capacity to process binaural cues, including interaural time differences (ITD) and interaural level differences (ILD). Our brain can determine differences in both arrival time and sound amplitude of incoming sound at each ear with remarkable accuracy, and it uses both as cues for sound source direction. Importantly, in natural hearing, the physical acoustics of the head and outer ears ensures that ITDs and ILDs are tightly coupled, such that, for example, sounds arriving from the left will arrive slightly earlier, as well as being slightly louder, in the left ear. However, in technologically assisted hearing, when stimuli are delivered to each ear separately by independent headphones or bilateral cochlear implants (CIs), ITDs and ILDs may become uncoupled. In the music industry, the production of stereophonic music tracks typically involves imposing ILDs on the recording of individual instruments or voices to place them to one side, but they leave the ITD at zero. When computing a sound source direction estimate, our auditory system combines information from both ITDs and ILDs, and when ITD and ILD cues disagree, we typically perceive the sound source at a “compromise location”, a phenomenon described as “cue trading” [1–5].

Cue trading is a sensible strategy when both types of cues are usually reliable, as is the case in normal acoustic hearing, and it is not particularly problematic in stereophonic music recordings, where zero ITDs pointing towards the midline at least do not completely contradict ILDs that place the guitar on the right and the singer on the left, so that the desired effect is easily achieved. The situation in prosthetic hearing with CIs is, however, quite different, because contemporary CIs will place separate devices at each of the listeners ear, each of which produce pulsatile electric stimulation, resulting in pulse timing ITDs that vary entirely randomly over a very wide range in a manner that is entirely uninformative about sound source direction and entirely uncoupled from ILDs. In this situation, ITDs may become not just uninformative, but positively misleading [6]. Interestingly, patients fitted with bilateral CIs typically exhibit poor sensitivity to ITDs [7; 8], but their ILD sensitivity tends to be good [9–11]. What causes the poor ITD sensitivity on bilateral CI patients remains somewhat controversial, and several researchers have favored the idea that a failure of normal binaural development occasioned by early onset hearing loss may be at least partly to blame. However, in light of the potentially disruptive influence of the randomized pulse timing ITDs delivered by clinical devices, it seems just as plausible to us that ITD sensitivity may be initially good in CI patients, as already shown in rats [12], but may subsequently be lost as the brain tries to adapt to the fact that the pulse timing ITDs delivered via clinical devices may be grossly misleading [6]. However, it is difficult to investigate this possibility in human volunteers because these volunteers usually only come to the tests after a long period of experience with their critical devices, and with unnaturally elevated ITD thresholds.

We therefore recently turned to the use of neonatally deafened, adult bilaterally implanted CI rats as an animal model, and we were able to show that these animals are consistently capable of discriminating even very small ITDs of 50-80 µs with only modest amounts of training, typically within a couple of weeks after activation of the CIs [12–14]. Furthermore, we recently showed that the median ILD sensitivity of neonatally deafened CI rats is at 1.7 dB [15]. Using this animal model, we have now been able to investigate how ITDs and ILDs interact in CI stimulation in a freshly implanted auditory system that has not undergone months of conditioning with randomized pulse timing ITDs. Our results, reported below, provide important hints that the randomized ITDs delivered by clinical CI stimulation do indeed have the potential to dramatically perturb ILD based sound source lateralization judgments.

Interactions between ITDs and ILDs are typically quantified by the “time-intensity trading ratio” (TITR), a psychophysical measure that indicates how many µs of ITD pointing in one direction are required to cancel out a 1 dB ILD pointing in the opposite direction [1; 3-5]. The TITR thus quantifies the relative strength of ITD and ILD cues on overall spatial perception in µs/dB. In subjects with normal hearing, TITRs are somewhat stimulus dependent, with both sound frequency [3; 16; 17] and sound intensity [5] influencing TITR values. TITRs reported in the literature for acoustic stimulation range from ∼2.5 to 185 µs/dB [3-5; 18-20], but how well these values predict what might be expected in the CI stimulated auditory system is unclear given, for example, that the encoding of acoustic stimulus intensity by the inner ear differs profoundly and in many ways from the encoding of electrical stimulus amplitudes at the electrode / auditory nerve interface.

## Results

Nine neonatally deafened (ND) rats were fitted with bilateral CIs in early adulthood (postnatal weeks 10-14). They were trained on a 2-alternative-forced choice (2AFC) sound lateralization task. A schematic representation of the behavioral training is shown in Figure 1 A. All rats were taught to initiate a trial by licking a center spout which triggered a 5 s long binaural biphasic pulse train of 900 pps presented directly through their CIs. The presentation of an ITD, ILD or both indicated to the rat which of two response spouts on either side of the central reward spout they had to lick to receive water as a reward. To determine the rats’ joint ITD and ILD sensitivity, we tested their responses to combinations of ITDs drawn independently from the set {0, ±60, ±80} µs and ILDs drawn from the set {0, ±1, ±4} dB. All possible combinations were tested, except (0 µs, 0 dB). We refer to trials in which ITDs and ILDs did not conflict (i.e. the leading ear is also louder or one of the cues was zero) as “honesty trials”, and in those trials the animals had to respond on the side indicated by both cues to obtain a reward. Trials where ITDs and ILDs were incongruent (i.e. the leading ear is quieter) we refer to as “probe trials”, and for these there is no objectively correct answer. Rather, the side that the stimulus is perceived on depends not just on the stimulus parameter values, but also on how each individual animal weights these cues, respectively. Given that the objective of the experiment was to measure this perceptual weighting without trying to shape it through reinforcement, all probe trials were rewarded irrespective of their choice. Figures 1 B and C show examples of honesty and a probe trial stimuli. To discourage animals from a strategy of random guessing without listening carefully to the stimuli, we randomly interleaved honesty and probe trials, with honesty trials outnumbering probe trials by approximately 4:1. See the Methods section for further details.

**Figure 1:**
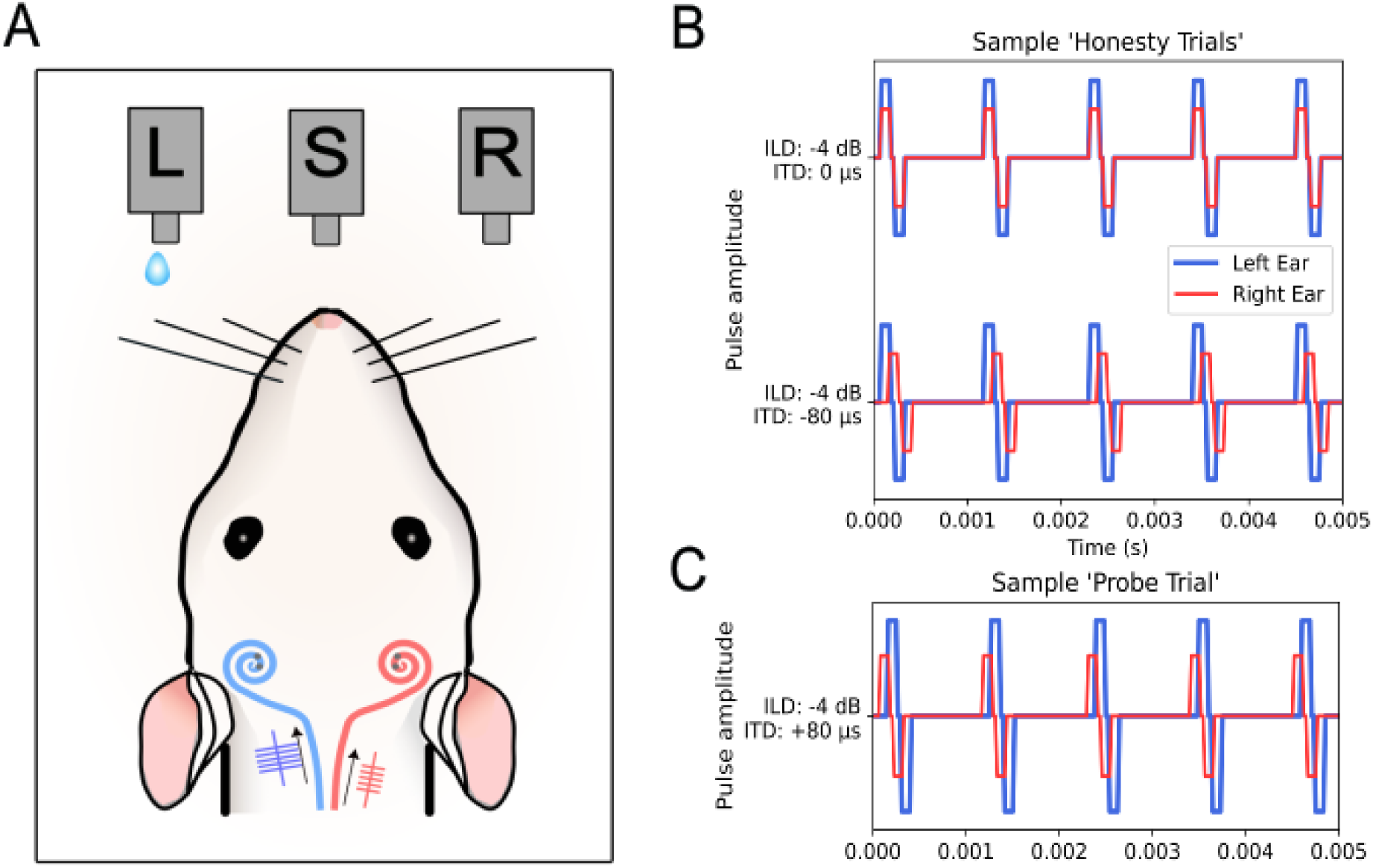
(A) Schematic of a neonatally deafened, cochlear implanted (CI)-supplied rat performing a two-alternative forced choice sound lateralization task in our custom-build behavioral setup. The rat needs to trigger the center spout (S) to receive a stimulus and then respond either on the left (L) or right (R) spout, accordingly. In this example, the stimulus arrives earlier and with higher intensity on the left CI, and only a response on the left water spout (L) would be rewarded. Two sample ‘honesty trials’ are displayed in (B). A binaural electric stimulus pulse train with an ILD of −4 dB and an ITD of 0 µs is shown in the upper panel, a pulse train with an ILD of −4 dB and an ITD of −80 µs is shown in the lower panel. In both examples the rat would need to respond on the left spout to receive drinking water as positive reinforcement. An incorrect response would lead to a timeout. (C) Example of a ‘probe trial’, here the binaural electric stimulus pulse train contains an ILD of −4 dB, pointing to the left, and an ITD of +80 µs, pointing to the right. During ‘probe trials’ the rat had a free choice and would be rewarded irrespective of the side they chose.

The nine animals were each tested in two sessions per day, five sessions per week, to collect between 849 and 7098 trials over the course of between six and 23 sessions. The raw data of the lateralization performance, pooled over all sessions for each rat, as a function of stimulus ITD and ILD, are shown as heatmaps in Figure 2. In each panel, responses to different ILDs are arranged bottom to top, while ITDs are arranged left to right. In each cell two numbers are shown. The top left indicates the number of trials for which the rat responded on the right reward spout, while the bottom right number reports the total number of trials performed for the specific parameter combination. The corresponding percentage of ‘right’ responses is also represented by the color of each cell: the lighter the color, the lower the percentage of responses to the right hand side and the darker the color, the higher the percentage of responses to the right hand side (see color bar bottom of Figure 2). This allows us to visualize the effects of ILD and ITD as color gradients running top-to-bottom or left-to-right, respectively. Most animals show a very clear gradient in the diagonal direction from top left to bottom right, as would be expected if the animals are sensitive to both ITD and ILD, and these two cues have more or less additive effects on the animals’ lateralization judgments. For some animals (e.g. rat 7), the left-to-right color gradient is a lot more obvious than the top-to-bottom gradient, indicating that these animals were a lot more influenced by ITD than ILD for the parameter ranges tested.

**Figure 2.**
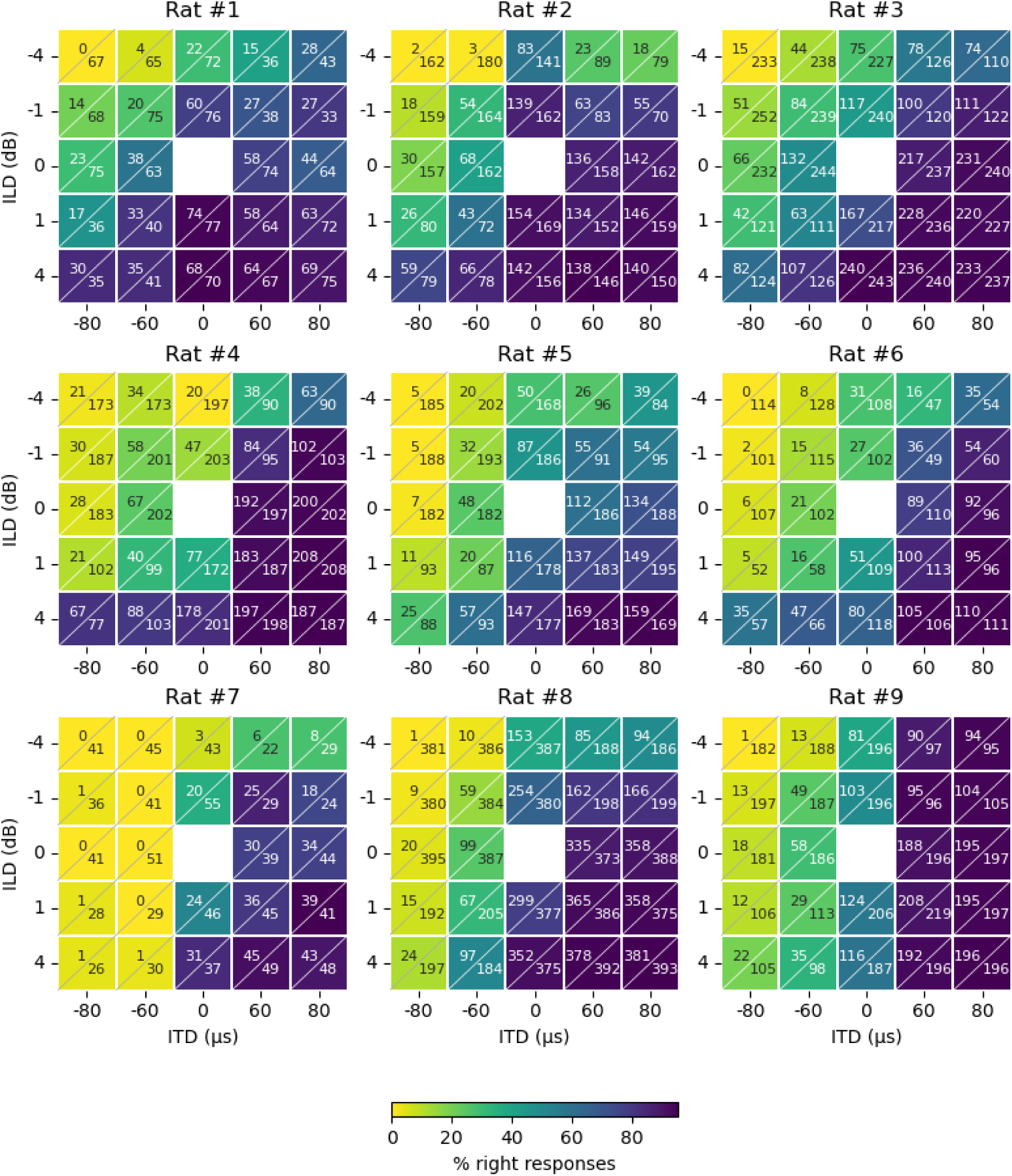
Heatmaps showing the proportion of responses to the right hand side for the different ITD (columns) and ILD (rows) combinations for nine neonatally deafened cochlear implanted rats (headings). Each cell shows two numbers; the upper left number indicates the number of responses to the right hand side and the lower, right number gives the total number of presented trials, resulting in the proportion of responses to the right hand side as indicated by the colorbar at the bottom.

To quantify the interaction of ITD and ILD on the binaural perception of our CI rats the data were fitted to a three-dimensional (3D) psychometric model. Figure 3 shows the fitted models in 3D plots for three representative examples. The fitted psychometric functions were adapted from the cumulative-gaussian-with-lapse functions we introduced in previous publications [13-15; 21]. The model assumes additive interactions between ITD and ILD cues and estimates sensitivity parameters for ITD and ILD, respectively (see Methods for details). Fitting psychometric functions to the data allowed us to estimate the “time-intensity trading ratios” (TITRs) which quantify the relative sensitivity to ITD and ILD as the ITD in μs pointing in one direction that would be required to cancel out a 1 dB ILD pointing in the opposite direction. Figure 3 shows examples of the fitted psychometric functions as mesh grids for three of the nine CI animals in our study. A strong ILD weighting is indicated by a steep slope of the mesh grid along the ILD-axis, while a strong ITD weighting is indicated by a steep slope along the ITD-axis. The examples shown in Figure 3 correspond to the animals with the largest (A), the median (B), and the smallest TITR (C) in our sample. Figure 3 D shows the distribution of TITR values across our cohort. The median TITR value across all nine CI animals was 18.7 μs/dB, the smallest was 3.9 µs/dB (rat 9) and the largest 27.1 µs/dB (rat 1).

**Figure 3.**
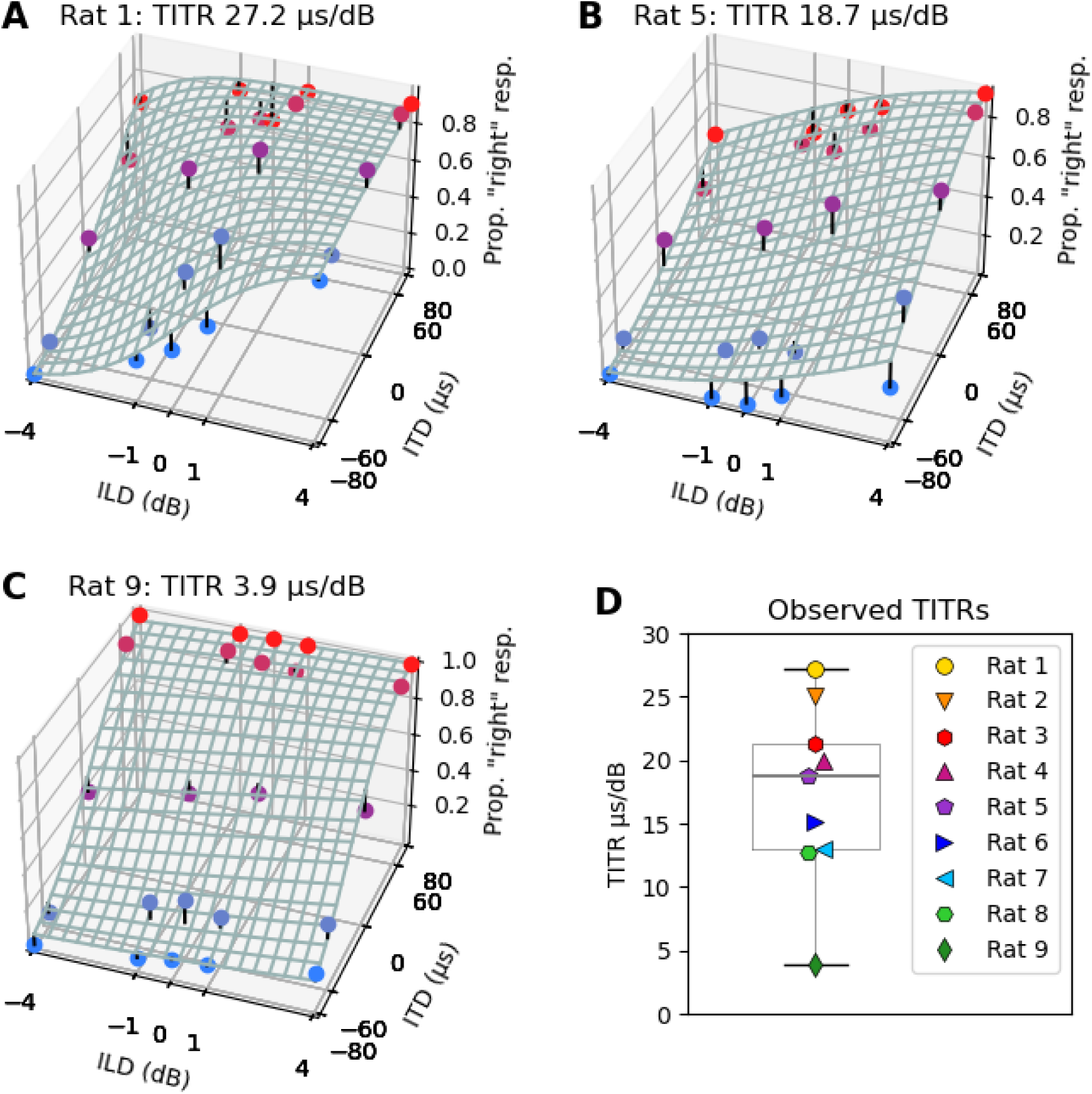
Sample three-dimensional joint ITD and ILD psychometric functions fitted to the raw data of three sample CI rats (A-C). X-axis: ILD values in dB with negative values representing higher intensity on the left ear. Z-axis: ITD values in µs with negative values representing ITDs arriving earlier on the left ear. Y-axis: proportion of trials for which the animal responded on the right hand side shown as colored dots in blue corresponding to left-leading ITDs and red to right-leading ITDs. The fitted psychometric function is shown as a mesh grid. The short black stems seen at some of the data points represent the residuals, that is the difference between the observed responses and those predicted by the fitted psychometric. Rats’ ID and individual time-intensity-trading-ratio (TITR) is indicated in the heading of each plot. (D) TITR values for each of the nine tested rats, indicated by differently colored symbols, and the median across all rats (18.7 µs/dB).

As an example of how the change of one binaural cue affects the overall binaural perception and the resulting lateralization decision, Figure 4 shows the percentages of responses to the right hand side side for two binaural cue combinations. When the combination of ILD = −4 dB (left leading) and ITD = 0 µs (frontal) was presented, the animals responded on the right side in on average 34 % of the trials. Adding an incongruent ITD cue of +80 µs (right leading) strongly affected the lateralization decisions of the rats and increased the percentage of responses to the right hand side by 25 %, resulting in a preference of the right hemisphere in 59 % of the trials.

**Figure 4.**
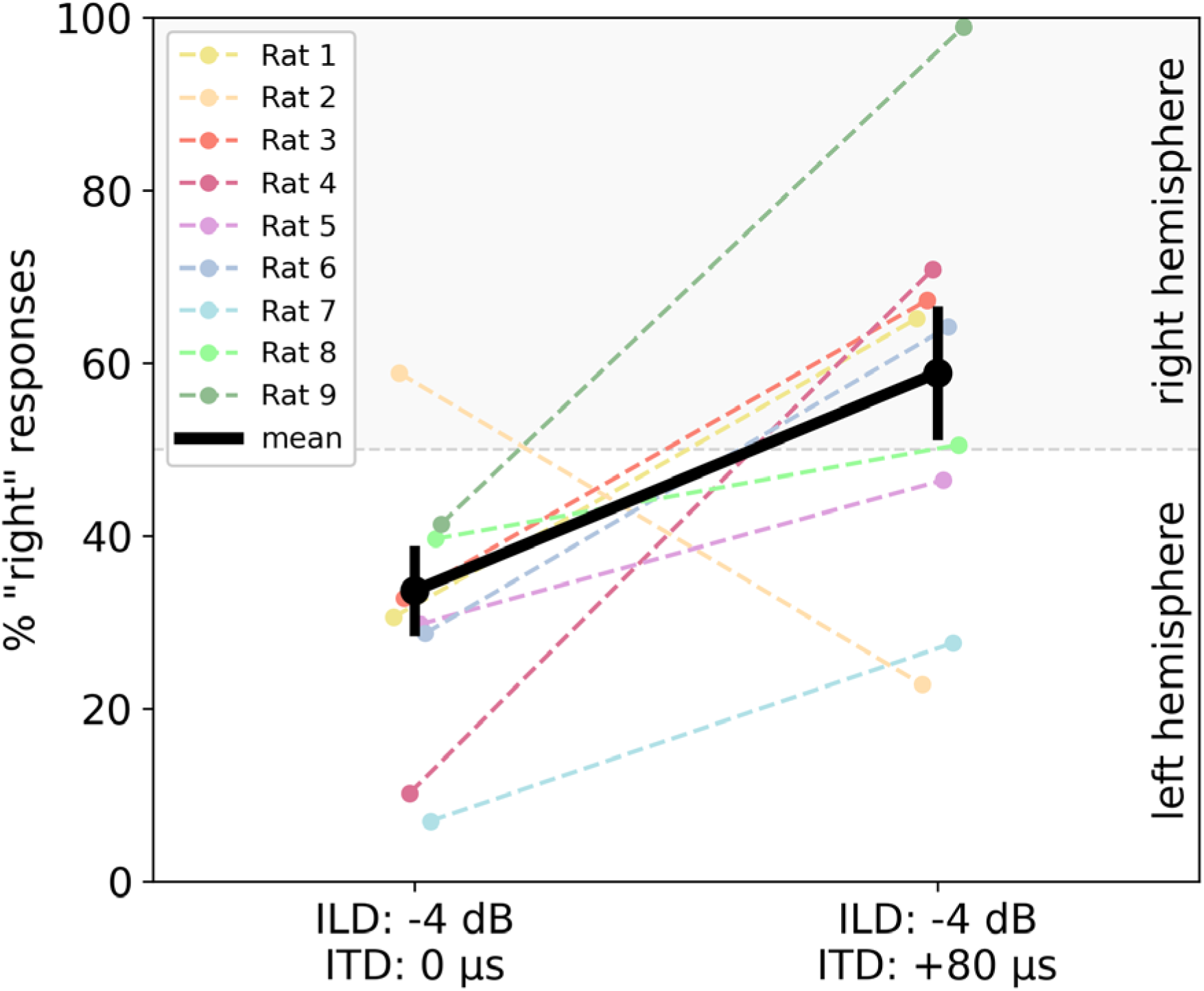
Percentage of responses to the right hand side of all nine CI rats for two binaural cue combinations: ILD −4 dB, ITD 0 μs and ILD −4 dB, ITD +80 μs. Zero to 50 % of responses to the right hand side correspond to the left hemisphere, while 50 to 100 % correspond to the right hemisphere. The individual CI rats are indicated by different colors, and the mean across all animals is displayed in black with error bars showing the standard error of mean.

## Discussion

The aim of this study was to investigate the interdependence and interaction between the two binaural cues ITD and ILD in electric hearing. To this end, behavioral experiments were conducted on neonatally deafened, adult CI-supplied rats, which have already proven to be an excellent model organism for binaural, electric hearing [13-15; 22]. Consequently, a TITR of only 18.7 µs/dB on average could be determined for the early deafened, electrically stimulated auditory system, which demonstrates not just the particular relevance of informative ITD cues for maximizing the potential for spatial hearing with bilateral CI devices, but also the large potential of inappropriate ITD cues to confound spatial hearing in situations where ILDs are reliable but ITDs are not.

### Binaural cue sensitivity in normal hearing and deaf mammals

Binaural cue sensitivity is generally well studied psychacoustically and electrophysically in the acoustic normal hearing system and, since the development of the CI, also in the deaf, electrically hearing auditory system. Acoustic ITD and ILD sensitivity is very comparable across mammalian species. ITD sensitivity, quantified by thresholds, so called just noticeable differences, is in the range of ∼6.9 to 96 µs, which can be found in detail in Table 1. Please note that the method used to determine the threshold may vary and that this is only intended as a rough guide.

**Table 1.** Interaural time difference (ITD) threshold across different normal hearing mammals.

| Species | ITD threshold | Source |
| --- | --- | --- |
| rats | ~50 μs | Li et al. 2019 [21] |
| cats | 30 μs | Wakeford and Robinson 1974 [23] |
| guinea pigs | 23-45 μs | Greene et al. 2018 [24] |
| rabbits | 50-60 μs | Ebert, Jr. et al. 2008 [25] |
| chinchillas | 55 μs | Koka et al. 2011 [26] |
| ferrets | 40 μs | Keating et al. 2013 [27] |
| gerbils | 12-96 μs | Tolnai et al. 2017 [28] |
| humans | 6.9-28 μs | Klumpp and Eady 1956, Zwislocki and Feldman 1956, Mills 1958, Thavam and Dietz 2019 [29-32] |

In comparison, the ILD sensitivity across different mammalian species is around 0.5-3 dB, details about specific thresholds can be found in Table 2. Please note that the compiled threshold values may vary in their measurement method and that this is intended only as a rough guide.

**Table 2.** Interaural level difference (ILD) thresholds across different normal hearing mammals.

| Species | ILD threshold | Source |
| --- | --- | --- |
| ferrets | 1.3 dB | Keating et al. 2013 [27] |
| guinea pigs | 3 dB | Greene et al. 2018 [24] |
| gerbils | 0.8 dB | Tolnai et al. 2017 [28] |
| rats | 3 dB | Wesolek et al. 2010 [33] |
| humans | 0.5-2.5 dB | Mills et al. 1960, Yost et al. 1988, Grothe et al. 2010, Laback et al. 2004 [9; 31; 34; 35] |

The ILD thresholds for human CI patients are in the range of 1-5 dB [9; 10; 36; 37], and thus show an ILD sensitivity comparable to NH listeners as well as CI rats (median ILD threshold =1.7 dB [15]). However, the picture is somewhat different with regard to ITD sensitivity. There are large differences between NH listeners and CI patients, with even the best candidates, irrespective of their hearing experience, not reaching NH thresholds [7; 29; 30; 32; 38-41]. In contrast, we have repeatedly shown that CI rats can achieve excellent ITD sensitivity with sub-millisecond thresholds even at clinical pulse rates up to 900 pps [13; 14; 22]. However, the type of stimulation seems to play a crucial role: while current clinical CI processors and stimulation strategies do not transmit microsecond precise temporal information, our custom-built CI stimulation setup for rats can provide microsecond precise ITD information. CI rats are thus a highly suitable model for investigating binaural cue sensitivity, with the unique opportunity to examine how the binaural cues ITD and ILD interact with each other.

### Weighting of ITD and ILD cues in acoustic and electric hearing

To date, the interplay of ITD and ILD cues has mainly be investigated in the acoustically stimulated auditory system, most likely because the overwhelming majority of bilateral CI patients do not have the requisite ITD sensitivity to make measurements of TITRs in such patients very meaningful. Our discovery that bilaterally CI implanted rats routinely exhibit excellent ITD sensitivity has changed this. In acoustic experiments, TITRs can be examined both electrophysiologically and psychophysically. For example, some studies have used single cell recording in the inferior colliculus of bats to evaluate TITRs. Mean TITR values varied between species from 47 µs/dB in Mexican free-tailed bats [42], 30.2+/-20 µs/dB in big brown bats [43], and as low as 17.9 µs in pallid bats, for which very similar values were also found on the level of the auditory cortex (16.7 µs/dB [44]). Psychoacoustic studies on normally hearing subjects have identified a number of stimulus parameters that can influence TITRs. For example, Gaskell and Henning (1980, [4]) found that the TITR for clicks and low frequency pure tones (∼50 µs/dB) was half as large as for high frequency amplitude modulated tones (100 µs/dB), suggesting that the stimulus waveform and frequency have an effect on the TITR. Additionally, adding background noise seems to have a negative influence on the ITD weighting for click and amplitude modulated stimuli. Furthermore, they showed that the TITR decreases with increasing ABL, i.e. one subject had a TITR of ∼13 µs/dB for the lowest intensity level of 48 dB, which decreased to less than 5 µs/dB for the highest intensity level of 93 dB. In addition, the ILD values used for the tests also appear to have an influence, with larger ILD values resulting in larger TITRs [5]. In addition to large individual differences, Stecker et al. (2010, [20]) found that the testing procedure can also make a difference. They tested normal hearing subjects with narrowband Gabor clicks in either a “closed-loop” or an “open-loop” procedure and found a significantly higher TITR for the “open-loop” (e.g. 80.2 µs/dB) than for the “closed-loop” procedure (e.g. 72.4 µs/dB). Furthermore, they examined the influence of the click rate and found a significantly higher TITR for both test procedures with decreasing inter-click interval from 10 ms (80.2 µs/dB; 72.4 µs/dB) to 2 ms (184.7 µs/dB; 118.7 µs/dB). Another important factor influencing TITRs is the frequency content of the acoustic stimuli used [3; 16-18]. To give just one example, Zerlin et al. (1969, [3]) have shown that the TITR increases with higher stimulus frequencies. At a center frequency of 1000 Hz, the TITR was 20 µs/dB, while at 4000 Hz the TITR reached 50 µs/dB.

In the framework of this study, using a behavioral testing paradigm, our CI rats reached TITRs ranging from 3.9 µs/dB (Figure 3 C) to 27.2 µs/dB (Figure 3 A), with a median of 18.3 µs/dB across all animals (Figure 3 B, D). Relative to the TITR values from acoustic experiments just reviewed, these values are at the low end, indicating a particularly pronounced influence of ITDs. Due to the differences between acoustic and electric hearing, direct comparisons are of course difficult. Nevertheless, the ITD weighting of our CI rats with in median 18.3 µs/dB may appear surprisingly strong, even when compared against the TITR of 25 µs/dB of normal hearing subjects at low frequencies (250 Hz) [3], where ITDs are generally thought to be the dominant cue. Based on the median of 18.7 µs/dB found in our CI rats, an ITD of only 80 µs would suffice to cancel out an ILD of 4 dB, as evidenced in Figure 4. This underscores that ITD has the potential to be a very powerful cue for CI users, provided that ITDs are delivered in an effective manner to their auditory system. However, current clinical CI processors are not well adapted to the task of providing microsecond temporal information. Clinical stimulation strategies typically deliver pulses at fixed rates of about 1000 pps, which makes it impossible to encode acoustic fine structure with precision. Furthermore, the clocks of the two processors at each ear run independently of each other and independently of incoming acoustic cues. As a consequence, the pulse timing ITDs delivered to the patients are effectively random numbers that could be as large as +/- 500 μs, large enough to cancel out the effect even of large ILDs. Given that, as we have shown, ITDs can be very powerful cues, it is to be expected that inappropriate, random ITDs delivered by contemporary clinical devices also have great potential to interfere with binaural hearing unless patients learn to become insensitive to this cue.

### The combination of ITD and ILD cues resulted in an additive effect

Our data not only provide insights into the effects of conflicting ITD and ILD, but also show how both cues interact when presented congruently. This can be seen in the diagonal color gradient in Figure 2, which is strong across all animals, indicating that the congruent presentation of ILD and ITD made lateralization easier leading to a more accurate lateralization performance. Sharma et al. (2023, [45]) made a similar observation for NH subjects, who showed a shorter reaction time when lateralizing clear directional, congruent ITD and ILD cues than when only one of the two cues provided directional information. They therefore concluded that two binaural cues are better than one. In fact, Klingel and Laback (2022, [46]) found a similar effect in CI patients. The linear regression curves fitted to the subjects’ lateralisation decisions exhibited a significantly steeper slope for the presentation of congruent ITD and ILD cues than for ILDs alone. This already indicated that the addition of a non-zero ITD to a non-zero ILD significantly increased lateralization at low pulse rates of 100 and 300 pps and could now be demonstrated/verified for the first time in our CI rats at a clinical pulse rate of 900 pps. The presentation of congruent ITDs and ILDs compared to the presentation of only one cue also affects the neurophysiological level.

### Training can affect the weighting of binaural cues

Previous literature indicates not only that TITRs can depend on a number of stimulus parameters and be subject to sizeable individual differences, it also indicates that the relative importance of different cues for spatial hearing may be influenced by experience, training, or even short term influences such as attention. The rats used in this study had a lot of experience lateralizing stimuli based on ITD cues alone (e.g. [22]) prior to the ITD/ILD joint sensitivity measurements described here, and it is possible that this may have favoured the low TITR values we observed. Ignaz et al. (2013, [47]) provide an example of short term plastic effects in cue weighting: they tested NH subjects on their ITD and ILD weighting, keeping either the ITD or the ILD constant, while they adjusted the other binaural cue until the stimulus was perceived centrally (0° azimuth). They found that, during the trading test, attention to the ‘to-be-adjusted’ cue appeared to increase, as did the weighting, indicating that the subjects weighted the cue they were able to manipulate more heavily. Meanwhile, Klingel et al. (2021, [48]) investigated whether the weighting of the binaural cues can be changed by training. They trained NH listeners in a sound lateralization task with additional visual cues reinforcing either ITD or ILD cues. In a following testing phase, they observed a reweighting of binaural cues and this within the first training session. A later study indicated that this type of cue reweighting is frequency dependent [17]. They tested and trained their subjects with different frequency bands and found that training in which ITD was reinforced only increased the ITD weighting for the low frequency band. In contrast ILD reinforcement resulted in a reweighting for mid and high frequency, but not low frequency, bands. Furthermore, the ability to reweight binaural cues has also been demonstrated in CI listeners. Klingel and Laback (2022, [46]) studied the reweighting of binaural cues in CI patients by subjecting them to visual reinforcement training. During training, the visual cue was congruent with the ITD but offset from the ILD cue. As a result, ITD weighting increased slightly but significantly for a pulse rate of 100 pps, while no training effect was observed for 300 pps. In contrast, the data of our CI rats show that a strong ITD weighting is also possible even at higher pulse rates of 900 pps, enabled by the presentation of informative pulse timing ITD and possibly facilitated by training.

## Conclusion

This study shows that the early deafened auditory system can develop excellent sensitivity to both small ITDs and small ILDs, and that, in the absence of clinical stimulation patterns with randomized pulse timing ITDs, it will tend to combine these cues in a manner that weights ITDs heavily, as evidenced by small, TITR values of only 18.3 µs/dB on average in our CI rats. This naturally high sensitivity of the CI stimulated auditory system to ITD cues suggests that generally poor ITD and generally better ILD sensitivity seen in the overwhelming majority of bilaterally implanted human CI patients may develop as an adaptation post-implantation as the patients’ auditory pathway attempts to reduce the potentially severe interference from random pulse timing ITDs. This interpretation of our finding thus provides an explanation for the poor ITD sensitivity observed in many CI patients, as well as pointing towards possible approaches to improve the binaural hearing outcomes of patients in the future by implementing informative pulse timing ITD-encoding in clinical stimulation strategies.

## Methods

All procedures involving experimental animals reported here were performed under license approved by the Regierungspräsidium (government council) Freiburg (#35-9185.81/G-17/124, #35-9185.81/G-22/067). We confirm that all of our methods were performed in accordance with the relevant guidelines and regulations and that our study is reported in accordance with the ARRIVE guidelines. A total of nine female Wistar rats were used in this study. All rats underwent neonatal deafening, acoustic and electric auditory brainstem response (ABR/eABR) recordings, bilateral cochlear implantation and behavioral training as described previously [13-15; 22] and briefly below.

### Neonatally deafening and cochlear implantation

All nine rats were neonatally deafened by receiving daily intraperitoneal (i.p.) injections of kanamycin from postnatal day 8–20 inclusively as described previously in Rosskothen-Kuhl and Illing 2012; Rauch et al. 2016; Rosskothen-Kuhl et al. 2018; 2024 [49–52]. The ototoxic effect of kanamycin is known to result in the destruction of inner and outer hair cells while the number of spiral ganglion cells remains comparable to that in untreated control rats [53; 54]. Severe to profound hearing loss (>80 dB) was confirmed by the loss of Preyer’s reflex [55] and the absence of ABRs to broadband click stimuli. Rats were raised to young adulthood (postnatal week 10-14) and then implanted simultaneously with bilateral cochlear implants (CIs) under ketamine (80 mg/kg) and xylazine (12 mg/kg) anesthesia. Two to three electrodes from either a PEIRA (animal arrays, Cochlear Ltd, Peira, Beerse, Belgium) or a MED-EL (3-patch animal arrays, Medical Electronics, Innsbruck, Austria) electrode array were inserted via a cochleostomy over the middle turn. The electrode leads were connected to a Hirose connector which was fixed on the vertex of the animal’s cranium with screws and dental acrylic to allow chronic electrical stimulation of the cochlea. The correct function of the CIs was confirmed using eABRs as described previously in [51; 52].

### Electric stimulation

The electric stimuli used to examine the animals’ eABR and behavioral spatial hearing sensitivity were generated using a Tucker-Davis Technology (TDT, Alachua, FL) IZ2MH programmable constant current stimulator at a sample rate of 48,828.125 Hz and delivered directly to the intracochlear electrodes of the head-mounted percutaneous connector as described above via a custom-made cable that was connected and disconnected before and after each training session. The cable included a slip-ring connector and was suspended above the animal’s head from a counterbalanced arm, which ensured that the animal could move freely in the behavior box without the cable becoming entangled or impeding the animal’s movement. One of the tip CI electrodes served as the stimulating electrode, the adjacent electrode as ground electrode. All electrical intracochlear stimulation used biphasic current pulses similar to those used in clinical devices (duty cycle: 40.96 µs positive, 40.96 µs at zero, 40.96 µs negative), with peak amplitudes of up to 300 μA, depending on eABR thresholds. For behavioral training, we stimulated all neonatally deafened, cochlear implanted (NDCI) rats ∼3.5-4.5 dB above these thresholds. Additionally, the average binaural level (ABL) at which the rats were trained and tested, were confirmed to be reliably detected, but not so high as to cause discomfort (rats will scratch their ears frequently, startle or show other signs of discomfort if stimuli are too intense) by careful observation of the animals’ behavior during spontaneous presentations of test stimuli. For full details on the electric stimuli and stimulation setup see Buck et al. (2023), Rosskothen-Kuhl et al. (2021), Schnupp et al. (2023), Buchholz et al. (20024) [14; 15; 22; 56].

### Psychoacoustic training and testing

After implantation, the rats were trained in a 2AFC sound lateralization task in our custom-built behavioral setup as described in Buck et al. (2023), Rosskothen-Kuhl et al. (2021), Schnupp et al. (2023), Buchholz et al. (20024) [14; 15; 22; 56]. The behavioral setup consists of a training cage with three water spouts on one side. A LED would indicate the start of the trial at which point the rats were trained to lick the center spout which would trigger a bilateral CI pulse train. The animals then had to choose between the left and right spout to indicate on which side they heard the stimulus. Correct responses were rewarded with drinking water, incorrect responses triggered negative feedback in the form of a short timeout.

Prior to testing, the rats took part in another behavioral study, which will be reported elsewhere. However, all rats were trained with 900 pps pulse trains containing co-varying ITDs ±{40, 60, 80, 100, 120} µs and ILDs ±{0.5, 1, 2, 3, 4, 5, 6} dB as spatial cues with negative values indicating a leading left ear and positive values a leading right ear. The pulse trains were up to 5 s long, but animals were free to make a choice as soon as they wished after stimulus onset, and the stimulus was terminated as soon as the animal touched one of the response spouts. On average, animals responded within approximately 2 s after stimulus onset.

After the initial training, which lasted between eight and 14 weeks, TITRs of the animals were determined by testing their responses to combinations of ITDs drawn independently from the set of {0, ±60, ±80} µs and ILDs drawn from the set of {0, ±1, ±4} dB. These values were presented to the animals both congruently and incongruently in all possible combinations, except (0 µs, 0 dB). The trials with congruent ITDs and ILDs, which could additionally contain either 0 dB or 0 µs, served as “honesty trials” for which the rats had to respond correctly to receive a reward. For “probe trials” where ITDs and ILDs were presented incongruently, the animals were always rewarded.

On average, 17 sessions per rat were included in the analysis, with approximately 200 trials per session. Overall, this resulted in an average of 4070 trials collected per animal, of which ∼82 % (3356) were honesty trials and ∼18 % (714) were probe trials. Only sessions in which the rats scored >75 % for honesty trials were included to ensure that the rat listened properly and decided according to its real perception during the probe trials.

### Data Analysis

To determine the behavioral ITD and ILD sensitivity of our CI rats, the proportion of responses to the right hand side (p_R_) was fit as a function of stimulus ITD and ILD using a psychometric function adapted from the cumulative-gaussian-with-lapse model described in our previous studies [14; 15; 21; 56], under the assumption that ITD and ILD have additive effects. The model takes the form:

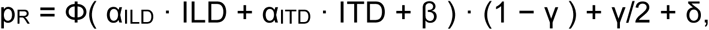

where Փ is the cumulative gaussian distribution function, ITD is the stimulus ITD in μs, ILD is the stimulus ILD in dB dB, α_ILD_ and α_ITD_ capture the animal’s sensitivity to ILD and ITD, and have units 1/dB and 1/μs, respectively, β allows for a possible “ear bias” (if stimuli with 0 ITD and 0 ILD are heard slightly off-center), δ captures a possible “spout bias” (i.e., when an animal guesses, it may have an idiosyncratic preference for one side), and γ is a lapse rate parameter that captures the proportion of times the rat makes errors due to inattention or exploratory behavior, irrespective of its ability to discriminate the stimulus. The model was fitted to the observed data using custom code that employed the python library function scipy.optimize.minimize() to implement a gradient descent that maximized the likelihood of the observed data given the set of fitted model parameters. Confirming the suitability of the model for our data, inspection of the residuals, the difference between the observed responses and those predicted by the fitted psychometric, showed no recurring pattern across animals (Figure S2). TITRs were then computed as the ratio of the sensitivity parameters α_ITD_/α_ILD_. The TITR is defined as the ITD in μs pointing in one direction that is required to offset a 1 dB ILD in the opposite direction, i.e. α_ILD_ · 1 −α_ITD_ · TITR = 0, which implies TITR = α_ILD_/α_ITD_, or the perception of the stimulus as 0° azimuth.

## Acknowledgements

The work leading to this publication was supported by the German Academic Exchange Service (DAAD) with funds from the German Federal Ministry of Education and Research (BMBF) and the People Programme (Marie Curie Actions) of the European Union’s Seventh Framework Programme (FP7/2007-2013) under REA grant agreement n° 605728 (P.R.I.M.E. – Postdoctoral Researchers International Mobility Experience), the Research Commission of the Medical Faculty of the Medical Center at the University of Freiburg, and the charity “Taube Kinder lernen hören e. V.“, MED-EL Medical Electronics, Innsbruck, Austria (Research Agreement PVFR2019/2), as well as grants from the Hong Kong General Research Fund (11100219, 11101020 & 11103823), the Shenzhen Science Technology and Innovation Committee (JCYJ20180307124024360) and the Martin Lee Centre for Innovations in Hearing Health at Macquarie University. Eighteen cochlear implant animal arrays and the cochlear implant images of the graphical abstract were kindly provided by MED-EL Medical Electronics, Innsbruck, Austria (Research Agreement PVFR2019/2). The authors would also like to thank Theresa Preyer and Henrike Budig for their assistance in collecting data.

## Author contributions

JS, NRK, and SB designed research; NRK and SB performed the surgeries; SB collected the data; JS and SB developed the analysis pipeline; SB, JS, and NRK analyzed the data; SB, NRK and JS wrote the paper.

## Data availability

All data as well as the analysis code used to generate all the figures and statistical results included in this manuscript are available from the corresponding author on reasonable request. All data generated or analyzed during this study are included in this published article.

## Supplementary Materials

**Figure S1.**
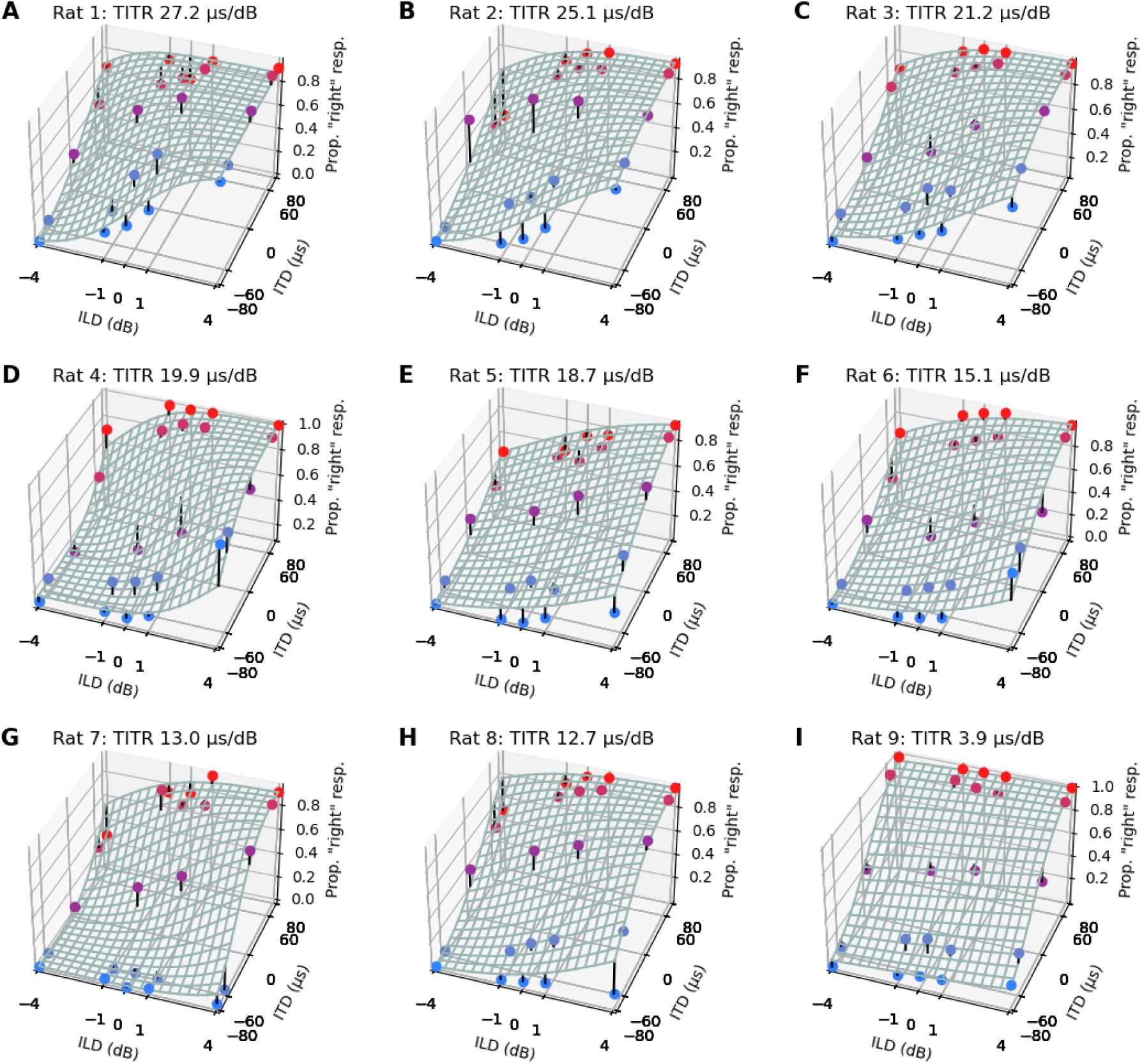
Three-dimensional joint interaural time difference (ITD) and interaural level difference (ILD) psychometric functions fitted to the raw data of all nine individual rats. X-axis: ILD values in dB with negative values representing higher intensity on the left ear. Z-axis: ITD values in µs with negative values representing ITDs arriving earlier on the left ear. Y-axis: proportion of trials for which the animal responded on the right hand side shown as colored dots in blue corresponding to left-leading ITDs, respectively. The fitted psychometric function is shown as a mesh grid. The short black stems seen at some of the data points represent the residuals, that is the difference between the observed responses and those predicted by the fitted psychometric. Rats’ ID and individual time-intensity-trading-ratio (TITR) is indicated in the heading of each plot.

**Figure S2.**
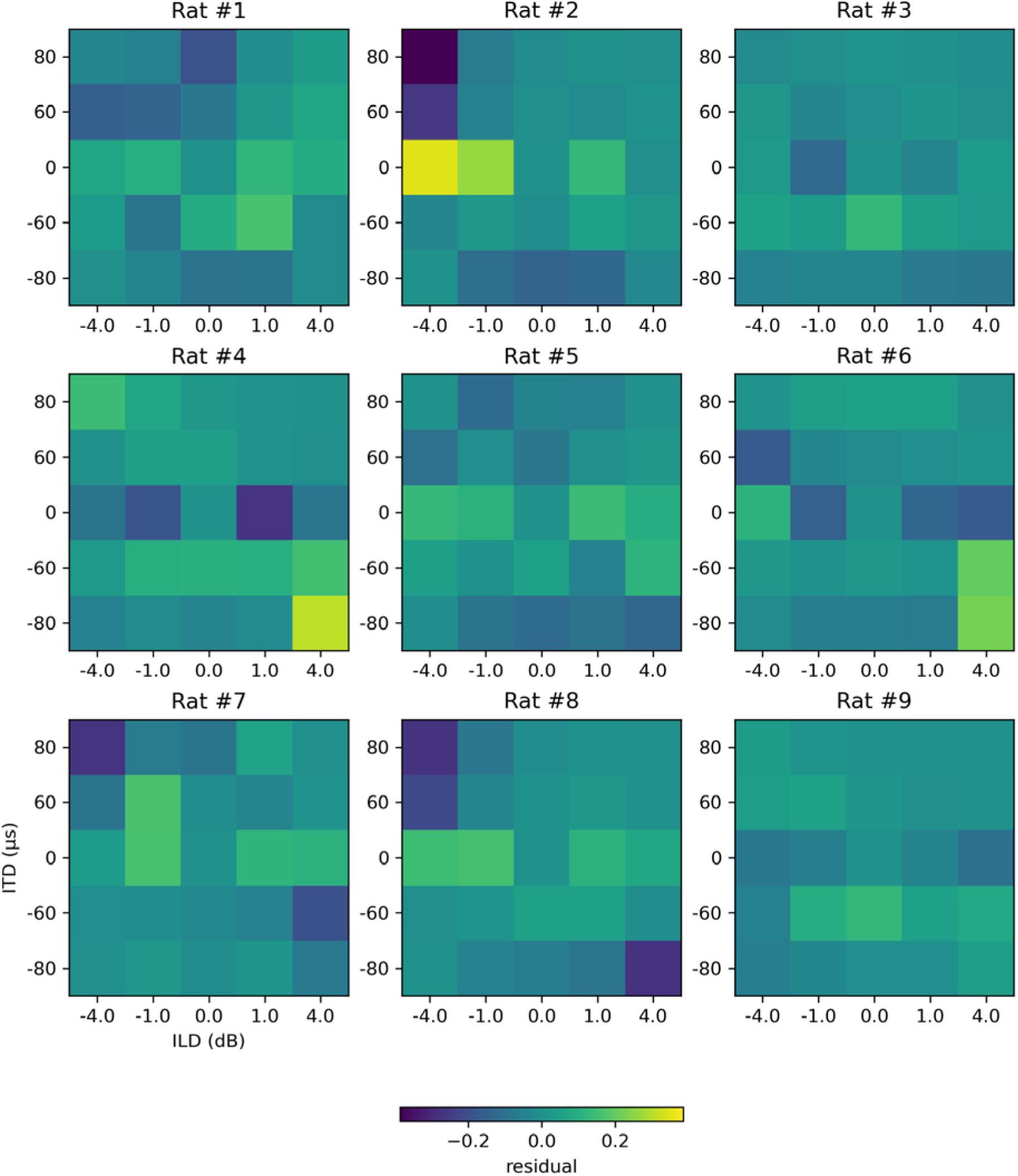
Heatmaps showing the residuals, which are the differences between the observed responses of our rats and those predicted by the fitted three dimensional psychometric, also visible as black stems in Fig. S1. The residual of each interaural time difference (ITD, columns) and interaural level difference (ILD, rows) combination is represented in colors according to their values as indicated by the colorbar.

